# Examining the Role of Notch Signaling in Dysplastic Lung Repair

**DOI:** 10.1101/2025.07.09.663914

**Authors:** Xinyuan Li, Madeline Singh, Andrew E. Vaughan

**Affiliations:** Department of Biomedical Sciences, School of Veterinary Medicine, University of Pennsylvania, Philadelphia, PA, 19104, USA; Institute for Regenerative Medicine, University of Pennsylvania, Philadelphia, PA, 19104, USA; Penn Lung Biology Institute, University of Pennsylvania, Philadelphia, PA, 19104, USA

## Abstract

Severe lung injury promotes the appearance of ectopic basal cells in the alveolar space. Since these cells appear to contribute to barrier restoration but demonstrate very inefficient differentiation into normal alveolar epithelium, this represents a dysplastic repair response. Prior work demonstrated the necessity of Notch signaling for expansion of these cells directly and indirectly via interactions with activated fibroblasts, but the original studies largely relied on γ-secretase inhibitors which lack specificity for the Notch pathway. Here we use transgenic mice expressing a dominant-negative MAML construct to confirm that Notch signaling to dysplastic cells impacts their expansion post-influenza. However, Notch inhibition at later time points did not appreciably increase differentiation into alveolar epithelial cells. These results confirm earlier studies and reiterate the advantages of genetic approaches to directly inhibit Notch over less specific pharmacological approaches.

## Introduction

Though critical for gas exchange, the lung is also a barrier organ and as such represents a common site of injury. In addition to exposure to environmental hazards and toxins, it is also subject to a wide range of respiratory pathogens. While many such pathogens can be effectively cleared while restricted to the upper airways, others, including some strains of influenza and coronaviruses, can infect distal epithelial cells and drive overexuberant inflammatory responses, causing severe injury to the blood-gas barrier and resulting in acute respiratory distress syndrome, which continues to bear a high mortality rate of >35%^1^.

In response to such severe injury, the mammalian lung employs a number of distinct progenitor cell types capable of responding to injury. In the airways, basal cells expressing the transcription factor p63 are well described to act as a progenitor cell for other airway cell type including club, ciliated, goblet, neuroendocrine, and tuft cells^2-7^. In the alveolar compartment, which is anatomically and functionally distinct from conducting airways, normally quiescent alveolar type 2 cells can transiently adopt progenitor features, proliferating and differentiating to replace alveolar type 1 cells responsible for gas exchange. However, in severe injury scenarios, this distinction between the airways and alveoli appears to be disrupted, in that basal-like cells can migrate from the airways into the alveolar region, aiding in barrier restoration but ultimately contributing only in very small part to the restoration of AT1s and AT2s^8-13^.

What’s more, these dysplastic regions of bronchiolized tissue appear to persist long term, at least 1 year in mouse models^14^. Identifying methods to promote transdifferentiation of basal cell-derived dysplastic tissue into more regionally appropriate alveolar cell types thus represents an important goal to aid in true, euplastic alveolar repair.

Several groups have identified signals involved in promoting dysplastic repair. For instance, HIF1a deletion or Wnt activation prior to injury blunts the dysplastic response in favor of greater contribution from distal airway secretory cells (club cells and BASCs)^9,15^ which appear quite capable of generating AT2s and AT1s^6,16^. Upon observing strong Notch target gene expression (Hes1) in dysplastic regions of tissue, initial work from our group demonstrated that treatment with γ-secretase inhibitors, widely used to inhibit Notch signaling via preventing cleavage of the intracellular domain, early in influenza injury similarly blunted Krt5+ cell expansion^8^. Follow-up work has now demonstrated that much of this effect can be ascribed to paracrine crosstalk between activated fibroblasts, with dysplastic basal cells providing Jag2 ligands to Notch3 receptors on the fibroblasts, which in turn generate trophic signals for basal cell expansion^17^. Treatment of post-influenza mice at later stages with γ-secretase inhibitors, in conjunction with dexamethasone and IBMX, resulted in a slight increase in AT2-like differentiation from Krt5-CreERT2 traced cells^8^. However, in spite of the widespread use of γ-secretase inhibitors to experimentally probe the Notch signaling pathway, it is increasingly recognized that γ-secretase has a very large number of substrates aside from just the Notch receptors, including but not limited to Eph receptors, Wnt receptors like Lrp6, E-Cadherin, and Trop2^18,19^. As such, it is difficult to ascribe any effect of γ-secretase inhibition solely to antagonism of the Notch pathway.

With this information in mind, here we reexamined previous models utilizing a genetically targeted approach to block all Notch signaling via inducible expression of a dominant negative MAML protein (dnMAML)^20,21^. This approach not only avoids the lack of specificity from γ-secretase inhibition but also has the advantage of reducing Notch signaling to the basal levels without the “derepression” phenomenon sometimes observed with RBPJ deletion^22^. As we recently reported, we observed that both dnMAML overexpression and treatment with the γ-secretase inhibitor DBZ significantly reduced expression of Notch target genes *Hes1, Hey1, and Hes2*, but only DBZ treatment exhibited significant growth and proliferation defects *in vitro* (see Supplemental Figure 12, Jones et al.^17^). We therefore utilized these animals to re-examine the role of Notch signaling in injury-induced basal cell expansion and differentiation *in vivo*.

## RESULTS

### Notch inhibition restricted to Krt5+ cells moderately but significantly inhibits dysplastic expansion after influenza infection

We recently reported that activation of dnMAML in injury-associated fibroblasts blocks further basal cell expansion and results in histologic improvement of tissue architecture after influenza infection^17^. We again confirmed that dysplastic basal cells highly express the classic Notch target gene Hes1 as compared to the surrounding alveolar parenchyma (Fig. 1A) after influenza infection. In order to quantitatively examine whether Notch signaling played any role directly in the Krt5+ cells, we induced dnMAML expression utilizing Krt5-CreERT2 mice at day 5 and day 7 post-infection, and harvested and homogenized whole lung tissue at day 11 (Fig. 1B). As we previously reported, we observed that induction of dnMAML expression directly in Krt5+ cells did not completely block their expansion. Recognizing the extreme heterogeneity of tissue injury after influenza infection, to avoid any sampling bias due to tissue sectioning and imaging analysis, we employed qRT-PCR analysis of whole lung RNA for both *Krt5* and *Tp63* transcripts. We observed a statistically significant ∼50% reduction in these surrogates for total dysplasia (Fig. 1B-C). Thus, while Krt5+ cells acting as a source of Notch ligand Jag2 for the alveolar fibroblasts appears to be the larger contributor to total dysplastic expansion, we confirmed that Notch signaling intrinsic to dysplastic Krt5+ cells does significantly contribute to their overall expansion after influenza-mediated injury^8^.

**Figure 1.**
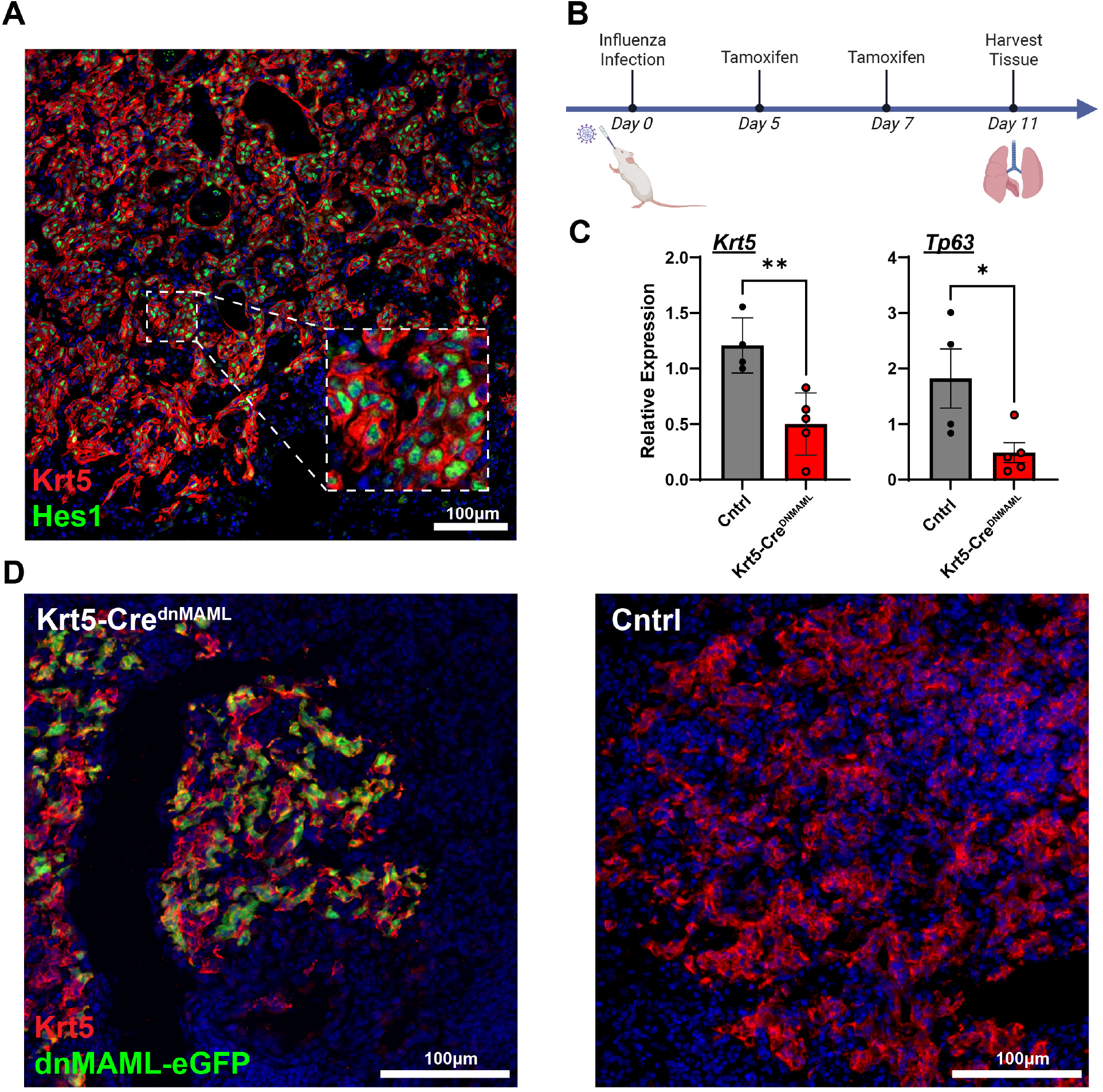
Krt5^+^ cell expansion is reduced following Notch inhibition after influenza injury. (A)Immunofluorescence staining shows nuclear localization and strong expression of Hes1 in Krt5^+^ cells at day 11 post-infection, indicating active Notch signaling. (B)Schematic timeline of the experimental design, including influenza virus infection, tamoxifen administration, and lung tissue collection for analysis. (C)Whole-lung RT-qPCR analysis at day 11 post-infection shows reduced expression of Krt5 and Tp63 in the Notch inhibition group compared to controls, n = 5 mice per group. (D)Immunofluorescence staining for Krt5 and GFP confirms dnMAML-mediated Notch inhibition in Krt5^+^ cells. The Krt5^+^ area is visibly reduced in Krt5-Cre^dnMAML^ lungs compared to controls.

### Inhibition of Notch signaling in later stage dysplastic repair

To address whether specific blockade of Notch in dysplastic cells at late time points impacted their ability to upregulate markers of alveolar type 2 cells, we utilized the above mice but also bred in a Cre-activated tdTomato allele to allow for lineage tracing as eGFP expression from the dnMAML construct tended to be weak. We allowed for full expansion of the Krt5+ cells and then administered tamoxifen to induce dnMAML expression at day 22 and day 25, alongside daily injections of dexamethasone and IBMX, and lungs were harvested at day 29-31. We observed very few cells expressing the canonical AT2 marker SPC in either condition (n = 5 mice per group examined) (Fig. 2). Noting important caveats (see Discussion), this generally argues against an important role for Notch signaling in regulating the rare fate conversion between ectopic alveolar basal cells and alveolar type 2 cells.

**Figure 2.**
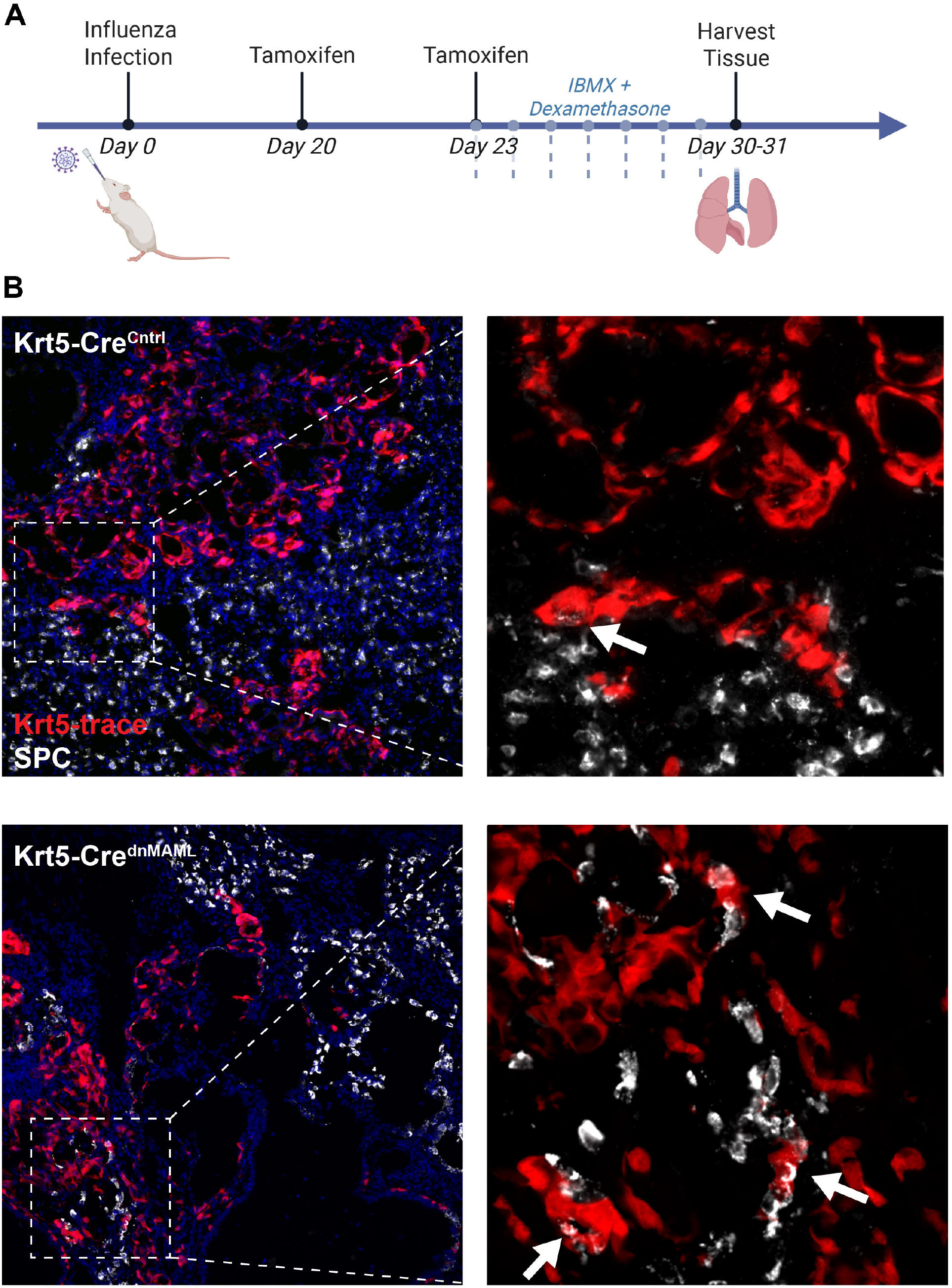
Genetic Notch inhibition in Krt5^+^ cells does not promote basal cell-to-AT2 differentiation. (A)Schematic timeline of the experimental design, including influenza virus infection, tamoxifen administration, IBMX and dexamethasone treatment, and lung tissue collection for analysis. (B)Immunofluorescence staining for Krt5 and SPC shows no obvious difference in AT2 cell differentiation between Krt5-Cre^dnMAML^ mice and controls.

## Discussion

Here, we utilized a well validated transgenic mouse model to reevaluate the role of Notch signaling in injury-induced basal cell expansion into the injured alveoli, often referred to as dysplastic repair. Recent collaborative work demonstrated that while Notch signaling is critical for dysplasia, Krt5+ cells act as a source of Notch ligands, signaling to adjacent alveolar fibroblasts, which in turn releases as-of-yet unidentified factors to promote Krt5+ cell expansion. Here, by blocking Notch signaling directly in the Krt5+ cells themselves and utilizing a highly quantitative method to measure total dysplastic expansion, we show that Notch signals received by the basal cells do still impact the overall expansion of injury-induced ectopic basal cells, albeit less than the effect of these cells sending Notch signals to adjacent mesenchyme as reported in our recent collaborative manuscript^17^. The source of the Notch ligands received by the Krt5+ cells themselves is uncertain, but since these cells exist as tight migratory clusters or sheets and express high levels of Jag2, we predict they largely provide these signals amongst themselves, to their own neighboring basal cells. These data largely reinforce prior work that in the context of injury, Notch signaling promotes basal cell proliferation even in their normal location in the upper airways^23^.

Notch signaling is well known to influence fate decisions between alveolar and airway cells in the developing lung. Several groups have demonstrated constitutive Notch signaling prevents alveolar specification in the developing lung, instead resulting in cystic structures lined by airway-like cells^24,25^. More recently, overexpression of dnMAML utilizing an airway club cell-specific CreERT2 followed by bleomycin injury demonstrated a nearly 30% increase in the ability of club cells to differentiate into AT2s upon Notch inhibition. Utilizing clever multicolor labeling and Notch intracellular domain (ICD) overexpression with the same club cell driver and bleomycin injury, it was demonstrated that nearly all Notch-active cells retained an airway club cell identity whereas labeled club cells without Notch activation greatly contributed to AT2 generation^26^. Another recent study indicated that this may be due specifically to Notch signaling dynamics in bronchoalveolar stem cells (BASCs)^16^. Similarly, in humans respiratory airway secretory cells (RASCs) are better able to generate AT2s upon γ-secretase inhibition^5^. Together these studies largely reinforce the paradigm observed in lung development wherein Notch promotes airway specification at the expense of alveolar cell types. Conversely, our results here show that Notch blockade specifically in ectopic basal cells after they arise had little effect on their conversion into SPC-expressing AT2s or AT2-like cells. That being said, there are important caveats to interpreting these results. We have noted that induction of dnMAML utilizing Krt5-CreERT2 causes severe toxicity apparent in the skin and possibly internal organs, making it difficult to keep animals alive after tamoxifen administration. Previous work with γ-secretase inhibition (DBZ) induced Krt5-CreERT2 lineage labeling at day 10 in order to label all Krt5 cells and their progeny, which ultimately includes numerous secretory cells as well as tuft cells and likely ciliated cells, and then administered DBZ, IBMX, and dexamethasone for 1 week starting 30 days post-influenza^8^. We were unable to induce lineage labeling and dnMAML expression at day 10 in this experiment, instead inducing recombination at ∼3 weeks post-infection. At this time point most basal cell expansion has already peaked and many of the daughter cells have already differentiated, thus avoiding labeling and dnMAML induction. As such, this experiment likely misses any contribution of Krt5+ cell-derived club cells that may more readily contribute to AT2s upon Notch inhibition.

Taken together, this work alongside recent publications provides important mechanistic and conceptual updates as to the role of Notch in the expansion and differentiation of ectopic, dysplastic basal cells in the lung after severe injury. While Notch signaling both within neighboring mesenchymal cells in the basal cells themselves promotes basal cell expansion, we do not observe strong evidence for a role of Notch cessation in promoting AT2 generation from these cells. This work also highlights that care needs to be taken when utilizing γ-secretase inhibitors and the recognition that many pathways other than Notch are modulated by these drugs. Given that the bulk of evidence argues that while Krt5+ cell expansion may help restore barrier function in the short term but impairs adaptive regeneration in the long term, future studies should focus on identification of additional pathways that may be modulated to improve the differentiation of ectopic basal cells into more regionally appropriate alveolar type 1 and type 2 cells.

## Methods

### Animals and treatment

8-to 12-week-old mice were used for all experiments with males and females in roughly equal proportions. Experimenters were not blinded to mouse age or sexes.

Ai14tdTomato (Gt(ROSA)26Sortm14(CAG-tdTomato)Hze) (Jax #:007914), Krt5CreERT2(PMID: 21983963), and Rosa26dnMAML-GFP (PMID: 16230473) mice have been previously described. All studies were approved by the University of Pennsylvania’s Institutional Animal Care and Use Committees and followed all NIH Office of Laboratory Animal Welfare regulations.

### Influenza virus infection

Influenza virus A/H1N1/PR/8 was administered intranasally at 25–45 TCID50 units of median tissue culture infectious dose to mice (20–25 g, 35 U; 25–30 g, 45 U) dissolved in 30 µl of PBS as we previously described^14^. Mice that lost 10% or more of their initial body weight by day 9 were deemed adequately infected and used for all experiments involving influenza infection.

### Lung histology

Lung tissues were fixed in 3.2% PFA for 1 h at room temperature, rinsed three times with PBS at room temperature, incubated in 30% sucrose overnight at 4 °C and transferred to 15% sucrose, 50% optimal cutting temperature (OCT) compound at room temperature for 2 h. Fixed tissues were transferred to an embedding mold filled with OCT compound, frozen on dry ice and stored at −80 °C.

### Immunofluorescence

Freshly dissected lungs were fixed, embedded in OCT compound, cut into 7-μm-thick cryosections. Slides were post-fixed another 5 min with 3.2% PFA and blocked in blocking buffer (1% BSA, 5% donkey serum and 0.1% Triton X-100 in PBS) for 1 h at room temperature. Next, slides were probed with primary antibodies and incubated overnight at 4 °C. The next day, the slides were washed and incubated with the fluorophore-conjugated secondary antibodies (typically Alexa Fluor conjugates; Life Sciences) at a 1:1000 dilution for 2 h. Lastly, slides were again washed, incubated with 1 µM DAPI for 5 min and mounted using ProLong Gold (Life Sciences, P36930). The following primary antibodies were used: Rabbit anti-SPC (1:2000, Millipore Sigma), Rabbit anti-Krt5 (1:500, BioLegend, clone Poly19055), Chicken anti-Krt5 (1:500, BioLegend, clone Poly9059), Rabbit anti-HES1 (1:500, Cell Signaling Technology, clone D6P2U, #11988), Goat anti-GFP (1:200, Rockland Immunochemicals, #600-101-215S).

### Whole-lung RT-qPCR

Lung tissue was harvested and immediately snap-frozen in liquid nitrogen. The frozen tissue was mechanically homogenized, and approximately 10 mg of tissue powder was added directly to TRIzol reagent for lysis. Total RNA was extracted using the Direct-zol RNA Miniprep Plus Kit (Zymo Research) according to the manufacturer’s instructions. cDNA was synthesized using the iScript Reverse Transcription Supermix (Bio-Rad).

Quantitative PCR (qPCR) was performed using PowerUp SYBR Green Master Mix (Applied Biosystems) on a QuantStudio 6 Real-Time PCR System (Thermo Fisher Scientific). Gene expression was normalized to the housekeeping gene *Rpl19* (“L19”) and expressed as fold change relative to the average expression level. The following primer sequences were used: *Krt5* forward 5′-TCTGCCATCACCCCATCTGT-3′ and reverse 5′-CCTCCGCCAGAACTGTAGGA-3′; *Tp63* forward 5′-CCTGGAAAACAATGCCCAGAC-3′ and reverse 5′-GAGGAGCCGTTCTGAATCTGC-3′; *Rpl19* forward 5′-ATGTATCACAGCCTGTACCTG-3′ and reverse 5′-TTCTTGGTCTCTTCCTCCTTG-3′.

### Statistics

All statistical calculations were performed using GraphPad Prism 10. Unpaired two-tailed Student’s t-tests were used to ascertain statistical significance between two groups. All experiments were performed on biological replicates unless stated otherwise. For details on statistical analyses, tests used, size of n, definition of significance, and summaries of statistical outputs, see corresponding figure legend and the Results section.

## Acknowledgments

This study was supported departmental startup funds from the Biomedical Sciences Department at the University of Pennsylvania to A.E.V.

## Author Contributions

X.L. and A.E.V. designed experiments, performed data acquisition and analysis, and prepared the manuscript. M.S. performed data acquisition and analysis. A.E.V. oversaw experimental design and manuscript preparation.

